# Fish load impacts biofilter microbial communities and nitrifier populations during establishment of freshwater home aquaria

**DOI:** 10.64898/2026.08.12.743087

**Authors:** Alexander K. Umbach, Natasha Szabolcs, Laura A. Sauder, Josh D. Neufeld

**Author notes:** Corresponding author: Department of Biology, University of Waterloo, 200 University Avenue West, Waterloo, Ontario, N2L 3G1, Canada.

## Abstract

Newly established freshwater aquaria rely on development of biofilter nitrifying populations to prevent ammonia and nitrite accumulation that can negatively impact fish health. Although initial fish loads impact water chemistry of new aquaria, little is known about the corresponding impact on microbial community succession within freshwater aquarium biofilters. To address this gap, fourteen home aquarium systems were established, stocked with a range of fish loads, and maintained for eight months. Aquaria were sampled regularly to monitor nitrogen species, microbial community composition (16S rRNA gene sequencing), and the abundance of nitrifiers (qPCR). Aquaria with higher fish loads developed microbial communities that were compositionally distinct from those with lower fish loads, and were dominated by *Pseudomonas*, *Rhodobacter*, and *Planctomycetes*. These patterns are consistent with increased nutrient availability supporting biofilm development, whereas lower fish loads may delay biofilm maturation. Increasing the number of fish in an aquarium significantly increased maximum ammonia and nitrite concentrations, although both were ultimately depleted within similar timeframes across treatments. Comammox *Nitrospira* were among the most abundant biofilter nitrifiers and were present in all biofilter samples regardless of fish load. Ammonia-oxidizing bacteria were detected at relatively low abundance but showed increases in relative abundance within high fish load aquarium filters. Ammonia-oxidizing archaea were below sequencing detection limits and detected only at low levels by qPCR, suggesting that their establishment in aquarium biofilters may require higher initial inoculation or longer timeframes. Overall, these results demonstrate that fish load shapes microbial community development in newly established aquarium biofilters, and that comammox *Nitrospira* dominate among nitrifiers during early biofilter establishment.

## Introduction

Ammonia and nitrite are metabolic waste products that are toxic to fish at concentrations as low as 0.1 mg/L (7.1 µM) [1]. An important contributor to ammonia and nitrite toxicity in home aquaria is fish load, which is defined as the number of fish per unit of volume. Guidelines for operating home aquaria often recommend gradually adding fish to a maximum load of “one inch of fish per gallon” [2–4] to avoid “new tank syndrome”, which describes toxicity associated with ammonia and nitrite accumulation [5, 6]. Additionally, guides encourage 2 to 4 weeks of “cycling” for an aquarium, which details the gradual addition of fish or ammonia to establish essential nitrifying populations [7–9]. Despite these recommendations, home aquarium systems vary considerably in ammonia loading depending on decisions of their respective operators, such as the size and types of fish, rate of fish addition in early stages of establishment, and the amount of food provided. Because home aquaria are effectively closed systems, new aquaria can accumulate ammonia and nitrite rapidly unless these are actively removed by water changes or passively removed by nitrification. Biofilters are essential aquarium components because they circulate water, trap and remove waste using sponge filters, and provide high surface area physical substrates (e.g., ceramic beads, sponges) that support development of biofilm and associated nitrifying microbial populations that oxidize ammonia and nitrite [10–14]. However, new aquarium biofilters lack established nitrifying populations capable of rapidly oxidizing ammonia and nitrite to nitrate, and little is understood about how fish loading influences primary succession of nitrifiers and their associated microbial communities.

Previous research investigating freshwater aquarium biofilter microbial communities has focused on established biofilters. Early investigations suggested that nitrite-oxidizing bacteria (NOB) and uncharacterized ammonia-oxidizing bacteria (AOB) were responsible for nitrification in freshwater aquaria [10, 11]. Ammonia-oxidizing archaea (AOA) were discovered almost a decade later [15] and were subsequently shown to be the dominant ammonia oxidizers in freshwater aquarium biofilters [12]. After the discovery of complete ammonia oxidizing “comammox” *Nitrospira* [16, 17], further research reported that comammox *Nitrospira* were also among dominant ammonia oxidizers within freshwater aquarium biofilters [13, 18–20]. An initial study of nitrifier succession in freshwater aquaria demonstrated that AOA were the dominant nitrifier and were temporally and spatially stable [14], but this work was performed prior to the discovery of comammox *Nitrospira*. Previous studies have contributed to an understanding of nitrifiers in established aquarium biofilters, but no studies have investigated early succession of nitrifiers within these systems since the discovery of novel comammox bacteria. Newly established aquarium biofilters are likely sensitive to mechanisms of primary succession and selective pressures, such as nutrient loading, may influence microbial community development [21, 22]. Ammonia oxidizers vary in physiological characteristics, including a range of substrate affinities, with AOB possessing a lower apparent affinity, and AOA and comammox *Nitrospira* having relatively high apparent affinities [23]. These differences in substrate affinity among nitrifying guilds are thought to drive niche specialization. Accordingly, the succession of nitrifiers in new aquarium biofilters is likely driven by initial ammonia and nutrient inputs. Investigating how fish load shapes nitrifying guild composition and overall microbial community succession in new aquarium biofilters will therefore provide insight into the ecological dynamics governing these organisms, which are essential to proper aquarium function.

This work investigates how variations in nutrient inputs, mediated through fish load, influence microbial community succession of new aquarium biofilters. It is expected that nitrifier population abundances will shift temporally in response to developing aquarium conditions, transitioning from high-ammonia, low-nitrate conditions in early aquaria that favor AOB to low-ammonia, high-nitrate conditions in established aquaria that favor comammox *Nitrospira* and AOA populations. An increased fish load is expected to influence the rate at which successional changes occur, and measuring these changes will provide insight into recommended cycling guidelines [7–9].

To characterize local nitrifier guild succession within freshwater aquaria in response to fish load, fourteen 10-gallon freshwater aquaria were established and maintained for eight months, with recirculating side-mounted filters and a range of zebrafish (*Danio rerio)* abundances. Water samples were collected regularly to quantify ammonia, nitrite, and nitrate concentrations, and ceramic beads from the biofilters were collected throughout the experiment for DNA extraction and microbial community analysis.

## Materials and Methods

### Experiment design and aquarium maintenance

To investigate how fish loading influences microbial community succession, fourteen 10-gallon freshwater aquaria were established and maintained for eight months, housed in the aquatics facility at the University of Waterloo. Of the fourteen aquaria, seven pairs were populated with 0, 1, 3, 6, 9, 12, or 15 zebrafish (*Danio rerio*). All aquaria were equipped with an AquaClear 30 Power Filter containing BioMax ceramic filter beads, a sponge, and granulated carbon (Fluval, Hagen Inc.), and a single artificial plant (Figure 1). The aquaria were treated with 10% hydrochloric acid and rinsed thoroughly with municipal tap water prior to set-up. Each aquarium was filled with tap water and treated with 10 mL of Big Al’s Multi-Purpose Water Conditioner (Big Al’s, Canada) to neutralize chlorine and chloramine. No additional chemical or biological supplements were added. Aquarium temperature and pH remained between 20-22°C and 8.0-9.0, respectively, throughout the experiment. Fish were fed New Life SPECTRUM Community Fish Formula at a quantity of two pellets (approximately 5 mg) per fish per day (i.e., 0, 5, 15, 30, 45, 60, 75 mg/day). Aquaria were exposed to 12-hour light/dark cycles and evaporation was offset with additions of municipal water treated with Big Al’s Multi-Purpose Water Conditioner as required. This study was approved by the Office of Research Ethics at the University of Waterloo (AUPP #11-23).

**Figure 1.**
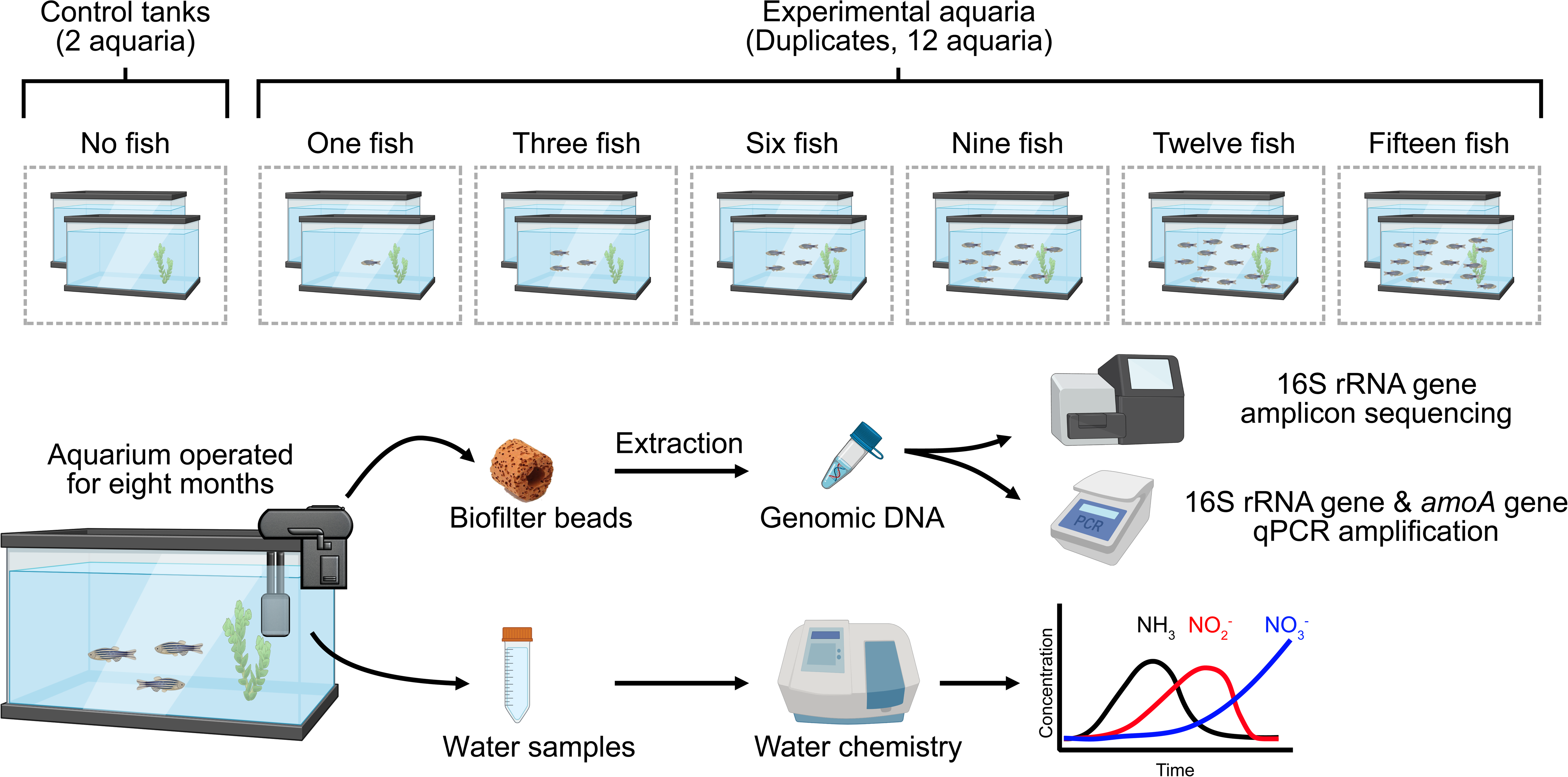
Schematic of experimental design and workflow for investigating the influence of fish load on aquarium biofilter microbial community succession. Made with BioRender. Alt text: Graphical representation of experimental treatments and replication and diagrams showing experimental workflows for data generation and analysis.

### Sample collection and processing

Ceramic biofilter beads were selected for sampling the microbial community because they contain similar communities compared to the sponge [24] but can be sampled without disturbing the entire biofilter. Ceramic beads were collected from each biofilter using ethanol-rinsed tweezers and stored in plastic tubes at -20°C until processing. Beads were collected once per week for the first six weeks, then once per month for the remainder of the experiment. Aquarium water samples were collected from each tank twice per week for the first four months, then once every two-to-three weeks for the remainder of the experiment and were stored in plastic tubes at -20°C until processing.

Genomic DNA was extracted from the ceramic biofilter beads using DNA Power Soil DNA Extraction kits (Qiagen, Germany), using a modified protocol to accommodate the biofilter bead (approximately 1 cm diameter). Specifically, the contents of the Power Bead tubes were transferred to a 5-mL conical tube along with 800 µL of the provided C1 lysis buffer. One biofilter bead was added to the tube and incubated at 70°C for 10 minutes in a rotating hybridization oven, followed by agitation on a benchtop vortex for 10 minutes at maximum speed. Tubes were then centrifuged at 7,000×g for 10 minutes, the supernatant transferred to a 1.5 mL tube, and the standard manufacturer protocol resumed.

### Quantification of ammonia, nitrite, and nitrate

Concentrations of total ammonia (NH_3_ and NH ^+^), nitrite (NO ^-^), and nitrate (NO ^-^) were measured for each water sample using an O-phalaldehyde (OPA)-based flurometric assay [25] and Griess reagent [26, 27] methods, respectively. Water samples were diluted 1:10 for ammonia and 1:5 for nitrite and nitrate. For the OPA method to measure total ammonia, 100 µL of diluted sample was combined with 100 µL of reagent in a black opaque 96-well plate and incubated in the dark at room temperature for four hours before measuring fluorescence at 360/465 nm using a spectrophotometer (FilterMax F5, Molecular Devices). For nitrite and nitrate measurements, 100 µL of diluted sample was combined with 100 µL of Griess reagent in a clear 96-well plate. An additional 100 µL of vanadium chloride solution (0.8 g VCl_3_ per 100 mL of 1M HCl) was added to the nitrate plate. Both plates were incubated in the dark at 37°C for one hour before measuring absorbance at 550 nm. Serial dilutions for ammonia (50 µM to 0.195 µM), nitrite (250 µM to 1.95 µM), and nitrate (500 µM to 3.91 µM) were used to construct standard curves to calculate relevant nitrogen species concentrations. Water chemistry data from replicate aquaria of all groups (i.e., 0, 1, 3, 6, 9, 12, 15 fish) were averaged for each time point.

### Amplicon sequencing and data processing

For microbial community profiling, genomic DNA was extracted from sampled ceramic beads (as described above), and the V4-V5 region of the 16S rRNA gene was amplified using universal prokaryotic primers 515F-Y (5’-GTGYCAGCMGCCGCGGTAA - 3’) [28] and 926R (5’ - CCGYCAATTYMTTTRAGTTT - 3’) [29]. Both primers were modified to include a 6-base barcode sequence used for identification of amplicons, an adaptor sequence for flow cell binding, and an Illumina primer binding site [30]. PCR amplification was performed in a sterile ISO 5 HEPA PCR hood, cleaned with 70% ethanol and treated with UV light for 15 minutes. A PCR master mix was created using UV-treated PCR-grade water, 1× ThermoPol buffer, 0.2 µM forward primer, 0.2 µM reverse primer, 200 µM dNTPs, 15 µg bovine serum albumin (BSA), 0.625 units of Hot Start *Taq* DNA polymerase (New England Biolabs, MA, USA), and 1 µL of DNA template in each 25-µL reaction. Amplification was performed using a T100 thermal cycler (Bio-Rad Laboratories, Canada) with the following conditions: 95°C initial denaturation for 3 minutes, 40 cycles of 95°C denaturation for 30 seconds, 55°C annealing for 30 seconds, 68°C extension for 1 minute, with a final extension at 68°C for 7 minutes. All PCR amplifications were performed in triplicate then pooled in equimolar quantities before purifying on a 1% agarose gel stained with ethidium bromide. Amplicons were extracted from the gel and purified using a Wizard SV Gel and PCR Clean-Up System (Promega, WI, USA). The library was diluted to 8 pM and 15% PhiX control v3 (Illumina, Canada) was added prior to sequencing. Samples were sequenced on a 2×250 cycle TG MiSeq Reagent Kit v2 (Illumina Canada, MS-103-1003) using a MiSeq System (Illumina) with the following quality: Q30 90.41, PF 92.82%, Cluster 526 ± 44 and PhiX measured at 25.3%.

Sequence reads were demultiplexed using the MiSeq Reporter software version 2.5.0.5 (Illumina). Demultiplexed sequences were processed using only the forward reads (single-end) to generate amplicon sequence variants (ASVs) and avoid exclusion of *Nitrospira-*associated sequences using QIIME2 2024.5 [31] and DADA2 [32]. Primers were removed using cutadapt [33] and forward reads were truncated to 224 bases and denoised using DADA2 [32]. The ASVs were classified using a naïve Bayes classifier trained against the SILVA 138 SSURef NR99 database [34, 35]. Negative controls were inspected manually to ensure that they were distinct from sample profiles. The ASV tables were then rarefied to 13,000 sequences using *qiime feature-table rarefy* and biological replicate samples for each time point were averaged using *qiime feature-table group*. Finally, ASV tables were filtered to remove ASVs with fewer than 10 reads to avoid analysis of spurious sequences generated during sequencing. All sequences generated for the current study were deposited in the European Nucleotide Archive (ENA) under project accession number PRJEB106317.

### Quantification of microbial communities using qPCR

To quantify AOA, AOB, and comammox *Nitrospira* populations, *amoA* and 16S rRNA genes were quantified in technical duplicate for each bead sample. Samples were diluted to DNA concentrations between 1 to 10 ng/µL and the *amoA* genes were amplified using primers CrenamoA23F/CrenamoA616R [36], amoA1F/amoA2R [37], and comaAFP/comaARP [38] for AOA, AOB, and comammox *Nitrospira*, respectively. Total bacterial and archaeal 16S rRNA genes were quantified using the universal primers 515F-Y/806R [28, 29]. Targets were amplified using 15 µL reaction volumes, constituted of 1X SSO SYBR qPCR master mix (7.5 µL of 2X stock) (Bio-Rad), 0.5 µg/µL BSA, and 2 µL genomic DNA (2 to 20 ng). Depending on the gene target, primers were added at 0.4 µM (AOA or AOB), 0.5 µM (comammox *Nitrospira*), or 0.3 µM (16S rRNA). Thermocycler conditions included an initial denaturation at 98°C for 3 minutes, followed by 35 cycles of 98°C for 30 seconds, annealing temperatures of 55°C for 30 seconds (AOA), 60°C for 30 seconds (AOB), 52°C for 45 seconds (comammox *Nitrospira*), or 50°C for 15 seconds (16S rRNA), followed by an extension temperature of 72°C for 60 seconds. Sample concentrations were calculated using standard curves of the *amoA* gene using representative sequences: *Ca.* N. aquarius (AOA, NCBI: KX034182.1), *N. europea* (AOB, NCBI: L08050.1:249-1079), *Nitrospira inopinata* (comammox *Nitrospira*, NCBI: MN836855.1), and *Thermus thermophilus* for the 16S rRNA gene.

## Results

### Total microbial community succession in response to fish load

To investigate microbial community succession, ceramic beads were collected throughout the experiment and genomic DNA was extracted for 16S rRNA gene amplicon sequencing. Sequence data revealed temporal changes in biofilter microbial community composition, as well as differences associated with fish load (Figure 2, Figure 3). During the first few weeks of operation, all aquaria were dominated by *Gammaproteobacteria* (>50% relative abundance), primarily attributed to *Thiobacillus, Pseudomonas*, and *Comamonadaceae*. In aquaria with fewer fish (i.e., n = 0, 1, 3), *Gammaproteobacteria* members remained relatively abundant throughout the experiment, although later time points increased relative abundances of ASVs affiliated with *Planctomycetota* (i.e., *Planctomycetes*, *OM190*), *Nitrospiria*, and low-abundance (<1%) taxa. In aquaria with higher fish loads (i.e., n ≥6), *Gammaproteobacteria* were displaced at later time points by greater relative abundances of taxa affiliated with *Planctomycetes* and *Nitrospiria*. Overall, *Planctomycetota* increased in relative abundance with both fish load and biofilter age.

**Figure 2.**
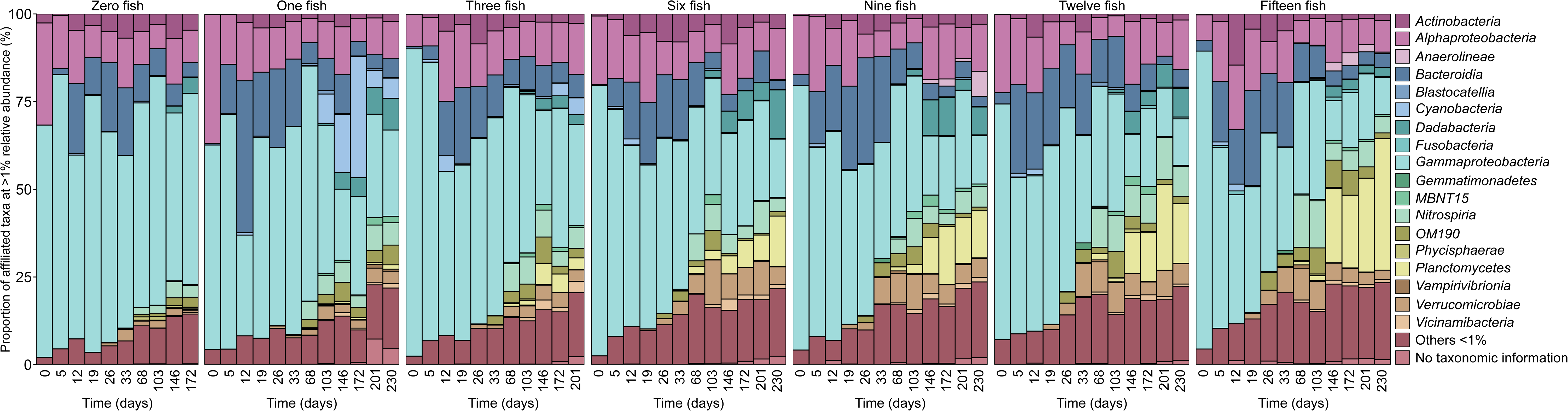
Class-level microbial community taxonomic distribution and relative abundance in aquarium biofilters. Taxa that were lower than 1% in any sample were grouped into the “Others” category. Beads for the “Zero fish” and “Three fish” aquaria were not collected beyond 172 and 201 days, respectively. Alt text: Stacked bar plots coloured by taxonomic class showing changes in relative abundances of taxa over time and among treatments.

**Figure 3.**
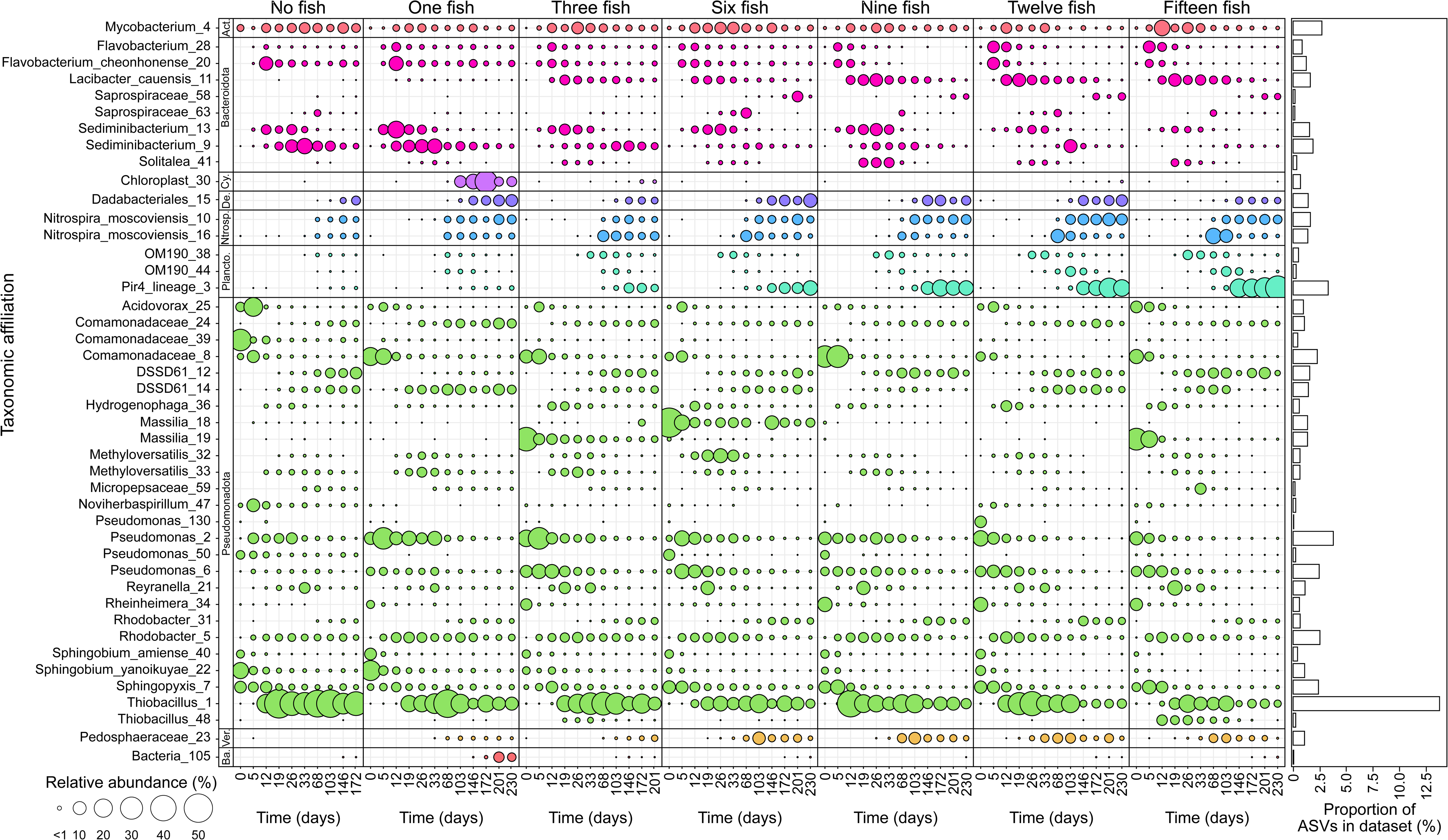
Distribution and relative abundance of microbial taxa among aquarium biofilters (16S rRNA). Bubble size represents the relative abundance of each taxon within a sample. Only taxa present at ≥ 5% in any sample are shown. Taxa are organized by phylum: *Actinomycetota* (Act.), *Cyanobacteriota* (Cy.), *Desulfobacterota* (De.), *Nitrospirota* (Nitrosp.), *Planctomycetota* (Plancto.), *Verrucomicrobia* (Ver.), and unclassified *Bacteria* (Ba.). Beads for the “Zero fish” and “Three fish” aquaria were not collected beyond 172 and 201 days, respectively. Alt text: Data displays showing relative abundance of amplicon sequence variants within each sample and the total proportions of each amplicon sequence variant within the dataset, organized by aquarium treatment.

Microbial community composition was further analyzed using both weighted and unweighted UniFrac metrics, with samples grouped into categories of “none” (n = 0), “low” (n =1, 3), “medium” (n = 6, 9), or “high” (n = 12, 15) fish load (Figure 4A and B). Early biofilter communities showed low dissimilarity across all treatments regardless of fish load, before differentiating temporally. Biofilter age (i.e., “Time”) was the primary predictor of community composition for both unweighted (*R*^2^ = 0.85, *p* < 0.05) and weighted (*R*^2^ = 0.83, *p <* 0.05) UniFrac metrics. Taxa associated with early time points included *Pseudomonas*, *Comamonadaceae*, and *Rhodobacteria*, whereas later timepoints were associated with *Thiobacillus* and a *Planctomycetes*-associated Pir4_lineage_3 ASV. Unweighted UniFrac analysis indicated convergence of microbial community profiles across all aquaria at later time points (days 201 and 230). In contrast, weighted UniFrac dissimilarities suggested separation of conditions by fish load, with communities from “none” and “low” fish load aquaria stabilizing earlier, whereas those from “medium” and “high” fish load aquaria converged only at later time points.

**Figure 4.**
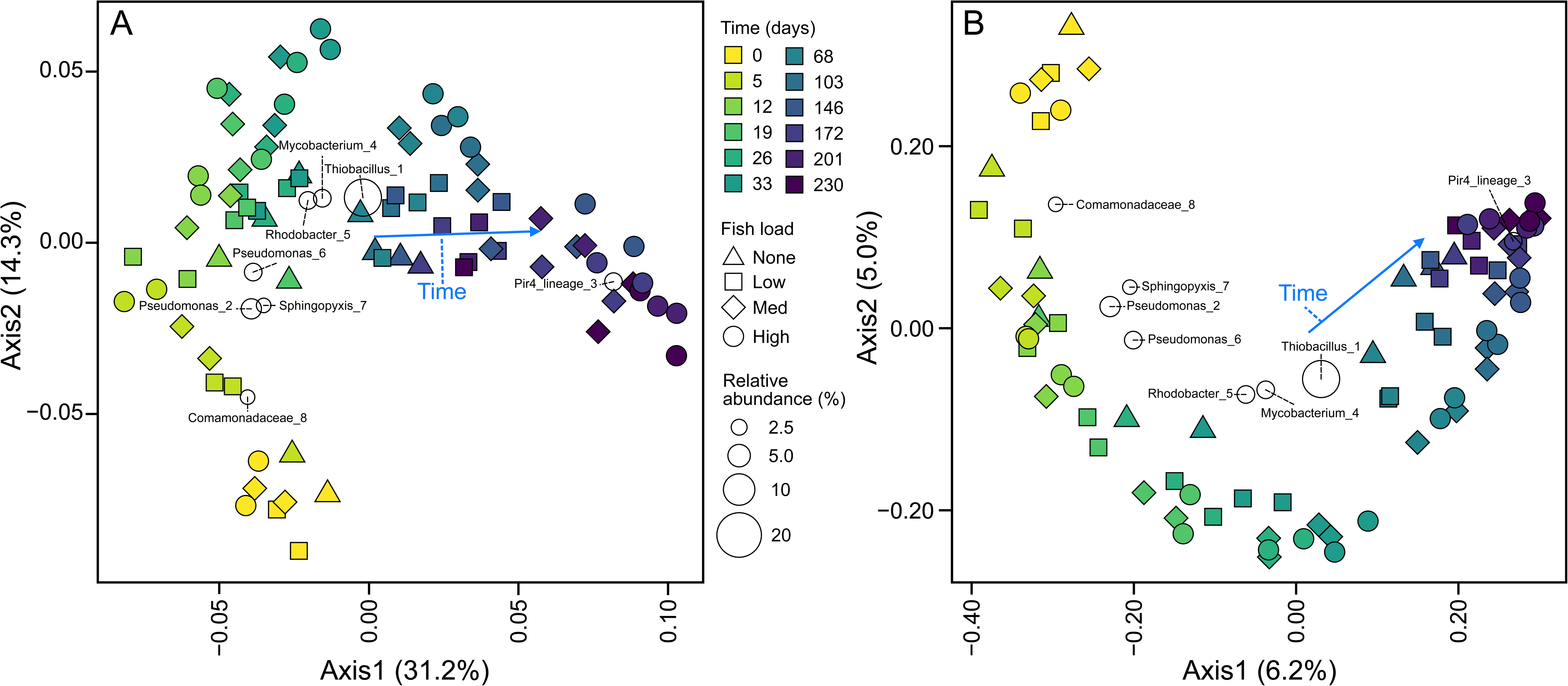
Microbial community composition and influential taxa using weighted (A) and unweighted (B) UniFrac. Only environmental variables with an *R*^2^ > 0.3 and *p* value < 0.05 are shown. Low, medium, and high fish loads are defined as aquaria containing one and three, six and nine, and twelve and fifteen fish, respectively. Alt text: Principal coordinate analysis plots showing how microbial community composition is influenced by both time and fish load.

### Nitrification and nitrifier guild succession in response to fish load

Water samples were collected throughout the experiment to monitor accumulation of ammonia, nitrite, and nitrate. As expected, increasing the abundance of fish resulted in increased ammonia and nitrite accumulation (Figure 5). Ammonia concentrations increased with fish load, with the highest peak observed in aquaria with the highest fish load (43.4 ± 3.4 µM, one-way ANOVA, *p* = <0.01). Nitrite accumulated during the first four weeks of operation, peaking between days 44 and 58 before decreasing rapidly. Similar to ammonia, peak nitrite concentrations differed among aquaria and were highest in aquaria containing more fish, reaching 214 ± 9 µM and 196 ± 87 µM in aquaria with 12 and 15 fish, respectively (one-way ANOVA, *p* = <1.6×10^-3^). Nitrate accumulated steadily throughout the experiment in all aquaria.

**Figure 5.**
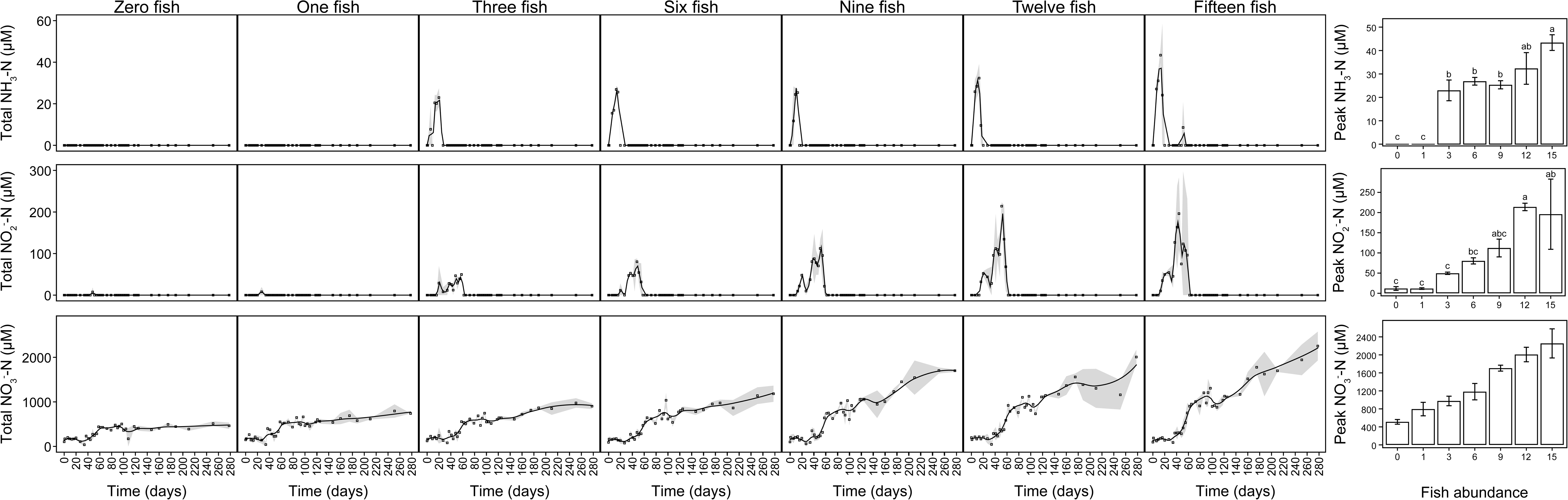
Ammonia, nitrite, and nitrate concentrations within experimental aquaria. Black squares represent average measured concentrations, and a black line modeled using a loess function shows the general trend. Grey shading represents the standard deviation of pooled technical and biological replicates. The bar plots to the right of the main panel displays the maximum detected concentrations for ammonia, nitrite, and nitrate. Letters above each bar indicate statistically significant differences. Alt text: Line plots and bar graphs depicting the rise and fall of ammonia and nitrite concentrations over time within each aquarium.

To investigate nitrifier guild and microbial community succession, 16S rRNA gene amplicon sequence data were combined with *amoA* qPCR data targeting specific nitrifying populations. For all aquarium biofilters, day 0 samples showed very low or undetected nitrifier relative abundances (Figure 6). Amplicon sequencing did not detect AOA-associated reads in any samples after data processing and quality control. Targeted qPCR for AOA *amoA* genes indicated low absolute abundances, with a trend of increasing AOA abundance with time for aquaria with higher fish loads (i.e., n = 12, 15).

**Figure 6.**
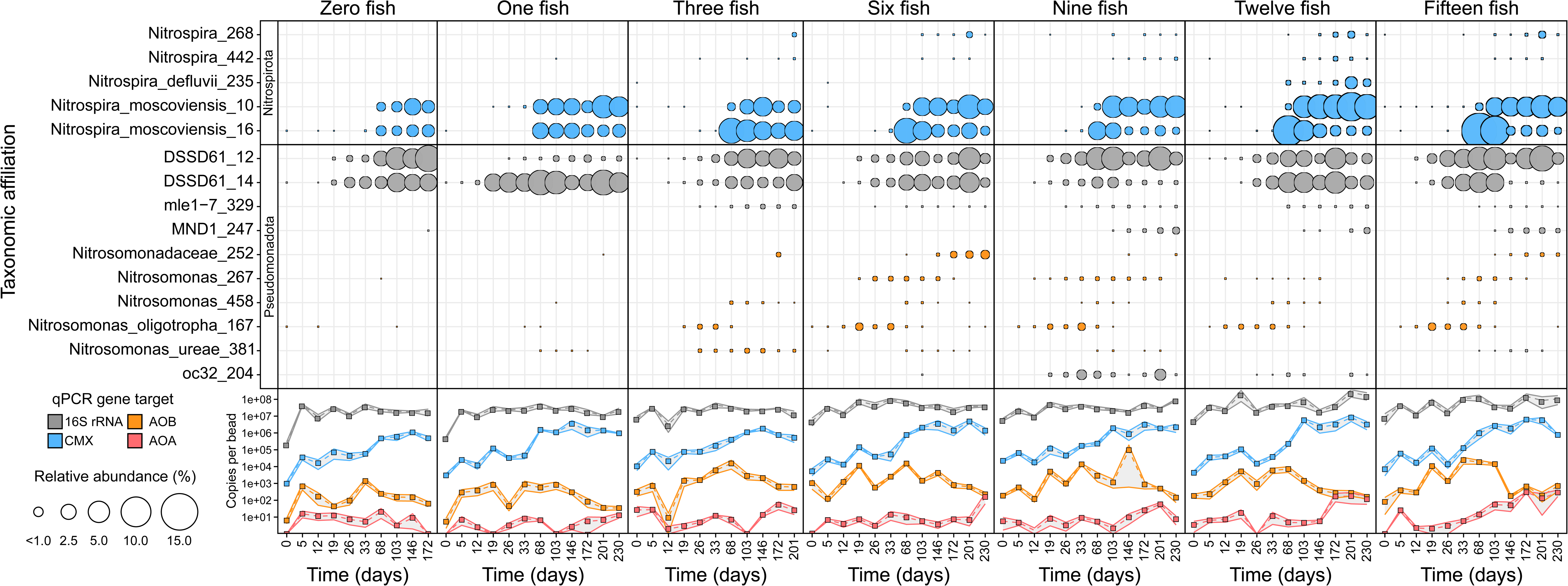
Distribution and abundance of putative nitrifiers within aquarium biofilters. Bubble size represents relative abundance of 16S rRNA genes within each sample. Line plots based on qPCR of the *amoA* and 16S rRNA genes show absolute abundances of AOA, AOB, and comammox *Nitrospira* (CMX) in relation to the total microbial community. Squares represent averages of biological and technical replicates and grey shading represents standard error of the mean. Beads for the “Zero fish” and “Three fish” aquaria were not collected beyond 172 and 201 days, respectively. Alt text: Graphs depicting both relative and absolute abundances of nitrifier-associated taxa within each aquarium, showing dominance of *Nitrospira* over time.

Absolute abundances of AOB measured by qPCR were low across all aquaria, with modest increases observed at intermediate timepoints (∼ days 26-103). Consistent with this finding, canonical AOB represented by *Nitrosomonas*-and *Nitrosomonadaceae*-associated ASVs were detected at low relative abundances in all samples. Several AOB-associated ASVs exhibited transient increases in relative abundance during early-to mid-timepoints before declining to low or undetectable levels at later stages. For example, *Nitrosomonas oligotropha*-associated ASVs were detected across all biofilters, increasing in relative abundance during the first ∼45 days before declining to undetectable levels. A second *Nitrosomonas*-associated ASV was largely absent from aquaria with lower fish loads but showed a similar transient pattern in higher-load systems. In contrast, a *Nitrosomonadaceae*-associated ASV was detected primarily at later time points and gradually increased in relative abundance.

Additional ASVs classified as *Nitrosomonadaceae* were detected at later timepoints, including members of the DSSD61 lineage and several low-abundance taxa (e.g., mle1-7, MND1, oc32). Several of these ASVs were observed at relatively high relative abundance across aquaria, particularly at later time points and in higher fish load conditions. However, these patterns were not consistent with AOB-targeted qPCR data, which indicated that AOB absolute abundances remained low throughout the experiment, suggesting that these taxa do not contribute to nitrification.

In contrast to AOB abundances, *amoA* gene copy numbers for comammox *Nitrospira* ranged between 9.6×10^2^ and 2.2×10^4^ copies per bead on day 0. Beyond day 0, amplicon sequence data indicate that *Nitrospira* populations were the dominant nitrifiers, increasing in both relative and absolute abundance over the duration of the experiment. Two core *Nitrospira* ASVs (both *Nitrospira moscovienesis*-associated) were prevalent for all aquarium samples, whereas additional *Nitrospira* ASVs (i.e., Nitrospira_268, 442, and Nitrospira_defluvii_235) were detected at lower and more variable relative abundances. Overall, these data indicate that nitrifying communities in aquarium biofilters were dominated by *Nitrospira*, with comparatively minor contributions from AOA and AOB.

## Discussion

### Biofilter microbial community succession

Amplicon data supported our expectations, demonstrating successional changes in microbial community composition that reflected differences in fish load and associated nutrient loading (Figure 2, Figure 3). Early aquarium microbial communities showed high similarity in the first week of sampling and were dominated by *Gammaproteobacteria* (*Pseudomonadales, Burkholdariales*) and *Alphaproteobacteria* (*Sphingomonadales*). These early communities are consistent with an initial heterotrophic bloom driven by dispersal [21] and colonization from surrounding environments. The initial time point likely represented microbial communities associated with ceramic beads, municipal water, and zebrafish microbiota, all of which can contribute common aquatic heterotrophs such as *Pseudomonas*, *Sphingomonadales*, *Comamonadaceae*, and *Acinetobacter* [39–45]. By the second week of operation, aquaria were exposed to influxes of organic carbon and nutrients through fish food and metabolic waste, introducing selective pressures that have potential to differentially impact microbial community succession. Sulfur-oxidizing *Thiobacillus* ASVs were particularly abundant throughout all stages of aquarium biofilter succession and were the most abundant taxa in the dataset, constituting 13.8% of all sequence reads (Figure 3). Sodium thiosulfate in the water conditioner used to treat the aquaria for chlorine and chloramine may have provided a competitive advantage to sulfur oxidizers in early low-nutrient aquaria, and throughout when the aquaria required additional water. Whether the sustained abundance of *Thiobacillus* reflects a transient response to sulfur inputs or a stable feature of aquarium biofilters remains unclear. This may also explain the abundance and prevalence of *Comamonadaceae*, given that they have also been reported to oxidize sulfur in addition to diverse organic compounds present within aquaria [46].

Microbial communities differentiated temporally, with aquaria containing more fish showing greater divergence from initial conditions and differentiation from aquaria with fewer fish (Figure 3). Members of the *Planctomycetes,* particularly *Pirellulaceae*-associated ASVs, increased in relative abundance temporally and primarily for samples associated with increased fish load aquaria. *Planctomycetes* members are commonly associated with freshwater biofilms and environments rich with organic carbon [47, 48] and have previously been detected in aquarium biofilters [13]. Their metabolism is diverse and includes simple and complex carbohydrate degradation [47, 49, 50] as well as nitrogen metabolism including anaerobic ammonia oxidation (anammox) [47, 51, 52]. It is not possible to identify whether anammox *Planctomycetes* are present in the system based on the chosen 16S rRNA gene region used for analysis, and a previous investigation showed little evidence to support anammox activity in large aquarium biofilters [53]. Although this does not preclude their presence, the observed increase in *Planctomycetes* more likely reflects their role as biofilm-associated heterotrophs supported by increasing organic carbon availability. Overall, the development of *Planctomycetes* populations is consistent with biofilm maturation in response to time and fish load.

Increased fish abundance leads to greater inputs of organic carbon and metabolic waste, supporting biofilm maturation and promoting growth of biofilm-associated taxa. Aquaria with higher fish loads reach biofilm maturation sooner than those with fewer fish, and aquaria with little or no fish may be incapable of supporting abundant biofilms, influencing overall microbial community composition. These trends are consistent with a transition from early colonization to more established, biofilm-associated communities influenced by nutrient loading. Indeed, a weighted UniFrac ordination showed separation of microbial communities at later time points, with aquaria under medium and high fish loads grouping separately from aquaria with fewer or no fish (Figure 4B). This was not observed as clearly with the unweighted UniFrac metric, indicating that late-stage biofilters contained similar taxa but that those taxa were differentially enriched depending on fish and nutrient loading.

### New aquarium nitrification and nitrifier populations

An important concern for hobbyists starting home aquarium systems is the toxicity associated with ammonia and nitrite accumulation. As expected, increasing the number of fish housed in the aquaria for this experiment was associated with increased ammonia concentrations (Figure 5). Ammonia was subsequently oxidized to nitrite, resulting in higher nitrite maxima in aquaria with a larger ammonia input. In general, the nitrifiers that mediate this transformation are nearly absent in new aquarium biofilters and the lag in associated nitrifying activity is the fundamental contributor to “new tank syndrome” and associated ammonia and nitrite toxicity [5, 6]. Indeed, after 1.5 weeks of aquarium operation, ammonia reached an average concentration of 0.6 mg/L in aquaria with the highest fish loads (Figure 5), a concentration potentially toxic to tropical fish [1]. Ammonia and nitrite were fully depleted after 60 days, which is to 3× to 6× longer than suggested for aquarium cycling [7, 8], and suggests that cycling periods exceeding the commonly recommended 2 to 4 weeks may reduce the risk of ammonia and nitrite accumulation.

Amplicon data implied that either AOB or comammox *Nitrospira* were present and contributing to early ammonia oxidation, whereas AOA were not detected following amplicon data processing and were only occasionally detected at low abundance by qPCR (Figure 6). The apparent ammonia affinities of AOB and comammox *Nitrospira* [23, 54] suggest selective pressure favouring high-affinity comammox *Nitrospira*, particularly under the relatively low ammonia concentrations observed in these aquaria. Absolute abundances of *amoA* gene copies support this interpretation, with comammox *Nitrospira* dominating the system. Short-read 16S rRNA gene amplicon sequencing data cannot distinguish between canonical NOB and comammox *Nitrospira* because of their recent evolutionary divergence [54, 55]. The dominant *Nitrospira* ASVs were taxonomically classified as “Nitrospira_moscoviensis”, which may traditionally indicate nitrite oxidizers rather than a complete ammonia oxidizer. However, given the high abundance of comammox *amoA* genes in the qPCR amplification data, this classification likely reflects limitations in taxonomic resolution, illustrating the challenge of identifying comammox *Nitrospira* populations using short-read 16S rRNA gene data. Together, these data suggest that comammox *Nitrospira* contributed the majority of ammonia oxidizing activity in these aquarium biofilters.

Sequences associated with AOB were detected throughout the experiment and increased in relative and absolute abundance until days 33 to 68, before declining thereafter (Figure 6). Because ammonia concentrations decreased to near or below detection limits (at ∼1 month), nitrifiers with high substrate affinity, such as comammox *Nitrospira,* may have outcompeted AOB that are better adapted for high-ammonia environments. This pattern is consistent with the persistence of large comammox populations and the corresponding decline of AOB. However, four ASVs taxonomically associated with *Nitrosomonadaceae* did not follow this trend, including DSSD61_12, DSSD61_14, mle1-7_329, and MND1-247. The two DSSD61 ASVs remained at relatively high relative abundance throughout the experiment, mirroring the succession of comammox *Nitrospira*. The mle-1 and MND1 ASVs were associated with aquaria containing three or more fish and were detected mostly in mid-and late-stage aquarium biofilter sample profiles. However, these patterns were not consistent with AOB-targeted qPCR data, which indicated low and declining AOB *amoA* copy numbers beyond day 68. This discrepancy may reflect incomplete primer coverage of diverse AOB lineages, given that the commonly used amo1A/2R primers were designed nearly 30 years ago based on cultured representatives of canonical AOB such as *Nitrosomonas europaea* [37]. Alternatively, these ASVs could be misclassified based on the 16S rRNA gene region used for phylogenetic and taxonomic analysis and may not represent ammonia oxidizers. If so, the associated *amoA* gene would be absent and thus explain the limited amplification within the qPCR amplification data.

Sequences associated with the DSSD61 lineage have only recently been reported in freshwater systems, and recent evidence suggests that these sequences may not belong to *Nitrosomonadaceae* [56–61]. In addition, there are no published genomes for DSSD61 representatives, and only a single associated 16S rRNA gene reference sequence (AY328759.1), isolated from a drinking water system, is currently available [39]. Nucleotide BLAST using both this reference sequence and ASV DSSD61_12 suggests that these sequences may instead belong to *Sterolibacteriaceae* or *Rhodocyclales* rather than *Nitrosomonadaceae*. Other freshwater aquarium surveys show that AOB are nearly absent or constitute a small proportion of the local nitrifying guild [7–9], suggesting that these taxonomically cryptic DSSD61 “AOB” are not nitrifiers. Considering the prevalence of short-read amplicon sequencing as the primary technique for characterizing microbial communities, the high relative abundance of misclassified taxa is problematic for studies generating conclusions regarding freshwater nitrifier ecology. Future work investigating taxonomic classification and metabolic potential of these cryptic DSSD61, OC32, MND1, and mle1-7 taxa is recommended.

Potential sources of nitrifiers include municipal drinking water systems and premise plumbing, which are known to contain populations of comammox *Nitrospira,* AOA, and AOB [62–69]. Although the municipal water used to fill the aquaria was not characterized, the high number of comammox *Nitrospira amoA* copy numbers at day 0 in all aquaria, including those with zero fish, suggests that these organisms were introduced from the municipal water supply during tank filling. The resident fish may also contribute to initial nitrifier seeding, as AOB have been detected in close association with the gills of teleost fish [70]. Despite low ammonia conditions that would be expected to favor AOA [12, 71], this group was largely absent from these aquarium systems. One possible explanation is that these experimental aquaria were established under controlled conditions with limited environmental inputs and maintained using clean techniques. In contrast, home aquaria operated by hobbyists are housed in common areas, are interacted with using bare hands and unsterilized equipment, and may include a number of added physical (e.g., soil, gravel) and biological (e.g., plants) materials through which AOA may be introduced. A recent aquarium survey detected members of the *Archaea* at relatively low abundance across several different systems and suggested that live plants may be a common vector through which AOA inoculate the system [13]. General features of the surrounding built environment may allow introduction of soil or debris containing AOA [72, 73]. It is possible that experimental aquaria maintained under home aquarium conditions, over several years, would result in biofilter nitrifying communities containing AOA as previously reported [12].

Given the variable and potentially low abundance of environmental sources of nitrifiers, aquarium owners often supplement new aquaria with commercial products containing live nitrifying cultures (e.g., Cycle, API Quick Start). Despite their popularity, there is limited research investigating their effectiveness. A single study showed that adding nitrifying supplements to “aquaria” containing only water and ammonia can influence nitrification rates [74], but this study lacked microbial profile data and the experimental design did not accurately reflect aquarium conditions. There are no known commercial nitrifying supplements that explicitly include AOA or comammox *Nitrospira*, which may constrain the effectiveness of these products in systems where high-affinity nitrifiers are favored. The efficacy of existing supplements should be investigated further, and developing new supplements that contain aquarium-relevant comammox *Nitrospira* and AOA could provide a more effective means to inoculate aquaria with beneficial nitrifiers and improve new aquarium outcomes. Inclusion of multiple nitrifying guilds may also increase robustness across a range of environmental conditions.

## Conclusion

Microbial community succession was influenced by fish load, reflecting differences in nutrient loading and biofilm formation. Biofilters were largely similar with respect to composition and succession of established nitrifiers. Comammox *Nitrospira* represented the dominant nitrifying populations in all aquarium biofilters after approximately 10 weeks, with AOB populations persisting at lower relative and absolute abundances in aquaria with three or more fish. Based on abundance data, nitrifying activity in all aquarium biofilters was primarily associated with comammox *Nitrospira*. The relative and absolute abundances of AOB were higher in aquaria with more fish than those with fewer, consistent with their adaptation to environments with higher ammonia concentrations. However, ambiguity surrounding the identity and function of several *Nitrosomonadaceae-*associated taxa highlights a need for additional investigation. The absence of AOA in amplicon sequencing data, together with their low abundance in qPCR, may suggest limited dispersion from environmental sources, that potential inoculation sources were lacking AOA, or that more time would be required to observe abundances consistent with previous aquarium biofilter studies. Together, these results provide a foundation for future research investigating nitrifier ecology in aquarium biofilters and support for the development of nitrifying supplements containing aquarium-relevant nitrifiers, including comammox *Nitrospira* and AOA, to improve performance and robustness across a range of environmental conditions.

